# An exploration of linkage fine-mapping on sequences from case-control studies

**DOI:** 10.1101/2021.12.18.473306

**Authors:** Payman Nickchi, Charith Karunarathna, Jinko Graham

**Affiliations:** Department of Statistics and Actuarial Science Simon Fraser University Burnaby, BC, Canada

**Keywords:** linkage analysis, fine-mapping, sequence relatedness, allelic heterogeneity, rare variants

## Abstract

Linkage analysis maps genetic loci for a heritable trait by identifying genomic regions with excess relatedness among individuals with similar trait values. Analysis may be conducted on related individuals from families, or on samples of unrelated individuals from a population. For allelically heterogeneous traits, population-based linkage analysis can be more powerful than genotypic-association analysis. Here, we focus on linkage analysis in a population sample, but use sequences rather than individuals as our unit of observation. Earlier investigations of sequence-based linkage mapping relied on known sequence relatedness, whereas we infer relatedness from the sequence data. We propose two ways to associate similarity in relatedness of sequences with similarity in their trait values and compare the resulting linkage methods to two genotypic-association methods. We also introduce a procedure to label case sequences as potential carriers or non-carriers of causal variants after an association has been found. This *post-hoc* labeling of case sequences is based on inferred relatedness to other case sequences. Our simulation results indicate that methods based on sequence-relatedness improve localization and perform as well as genotypic-association methods for detecting rare causal variants. Sequence-based linkage analysis therefore has potential to fine-map allelically heterogeneous disease traits.

## 1 Introduction

Linkage analysis is a classic tool to map genetic loci that contribute to a heritable trait. The basic idea is to look for genomic regions that have excess relatedness among individuals with similar trait values (Thompson, 2013). The approach therefore associates similarity in genetic relatedness with similarity in trait values (e.g. Almasy and Blangero, 1998; Amos, 1994). By contrast, genotypic-association analysis associates specific variants or aggregates of variants directly with trait values.

Linkage analysis has traditionally been conducted on related individuals from families. However, the use of families for fine mapping requires many informative meioses (Boehnke, 1994), either through numerous small pedigrees or large extended pedigrees, and enrolling such families may be impractical. An alternative that has been proposed for allelically heterogeneous traits is population-based linkage mapping (Purcell et al., 2007), which gains meioses by adapting linkage analysis to readily-available population-based case-control, cohort or cross-sectional samples. These methods scan individuals for excess ancestral sharing or identity by descent (IBD) at a locus, among individuals with similar trait values. The association between ancestral sharing and phenotypic similarity is assessed along the genome, and regions with high association are singled out for further study.

Browning and Thompson (2012) investigated the power of population-based linkage mapping to detect associations for complex diseases in case-control studies. They contrasted rates of IBD in case/case and non-case/case pairs of individuals at each single-nucleotide variant (SNV), and showed that IBD-based mapping has higher power than genotypic-association mapping when there are multiple, rare causal variants. Their results confirm the expectation that linkage analysis can be more powerful than genotypic-association methods for allelically heterogeneous traits (Ott et al., 2015).

The linkage analyses reviewed so far consider individuals as the unit of observation. Here, we take a different approach and use sequences as the unit of observation. We use sequences rather than individuals because, at a given genomic location, the gene genealogy connecting the sampled sequences groups them according to their relatedness. We assume that sequences which carry the same rare causal variant descend from a common ancestral sequence. As a result, they are IBD around the variant and will cluster together on its local gene genealogy. Genomic regions with excess trait clustering on their local genealogy therefore indicate a causal locus.

In this report, we explore the feasibility of linkage fine-mapping on sequences. In particular, we compare the ability of linkage and genotypic-association approaches to map an allelically heterogeneous disease of high penetrance. High-penetrance variants produce familial clusters that are easier to detect. As discussed in Ott et al. (2015), linkage methods are predicted to work well in such circumstances. We consider two linkage or descent-based methods that associate similarity in relatedness of sequences with similarity in trait values. For comparison, we consider two genotypic-association methods, one which considers single variants and another which aggregates variants. Through a simulation study, we compare the ability of these methods to fine-map rare causal variants in a 2 million base-pair (Mbp) candidate region. We chose 2 Mbp because it is the approximate resolution of a moderate-sized linkage study in pedigrees. For example, a linkage analysis with 100 informative meioses is expected to map a disease locus to within 2 centiMorgans (Boehnke, 1994), or approximately 2 Mbp.

To illustrate ideas, we work through an example dataset as a case study. Following this, we use a coalescent simulation to evaluate the ability of these methods to *detect* and *localize* a disease locus. Specifically, we are interested in the ability to detect any association within the candidate genomic region being fine-mapped, and also in the ability to localize the association signal to the causal subregion within the candidate region. Having detected a disease locus, it is of interest to identify case sequences that may be carriers of a causal variant. We conclude by describing a *post hoc* labeling procedure to classify case sequences into carriers and non-carriers of causal variants, using estimated sequence relatedness.

## 2 Materials and Methods

In this section, we describe how we simulated the genetic data and disease phenotype given the genetic data. Next, we describe the association methods that we considered to *detect* and *localize* causal variants. Finally, we propose a method for *post-hoc* labeling of case sequences into carriers and non-carriers of causal variants, given that an association has been detected.

### 2.1 Genetic-data simulation

We used *msprime* (Kelleher et al., 2016) to simulate the gene genealogy and sequences across a 2 Mbp genomic region for an entire population. We applied a hybrid strategy (D. Nelson et al., 2020) in which a backwards Wright-Fisher model with recombination and mutation was run to 5000 generations before present, followed by a coalescent with recombination and mutation from 5000 generations back to the overall most recent common ancestor across the genomic region. The hybrid strategy avoids inaccuracies in the coalescent approximation when the number of sampled sequences is large relative to the population effective size. The diploid population was of constant effective size, *N_e_* = 3100 (Tenesa et al., 2007), and consisted of 6200 sequences. We used a recombination rate of 1 × 10^−8^ per base per generation (Johnston and Cutler, 2012) and a mutation rate of 2 × 10^−8^ per base per generation (Campbell et al., 2012) to simulate 500 populations. Figure A.1 in Appendix A.1 displays the distribution of variant allele frequencies in the simulated population from which the example dataset was drawn. This allele-frequency spectrum is similar across the 500 simulated populations. The spike in the lowest-frequency bin of the histogram (frequency ≤ 0.01) is consistent with an observed abundance of rare variants in real populations; (e.g., M. R. Nelson et al., 2012).

### 2.2 Disease-trait model

To mimic random mating in a diploid population, we randomly paired the population sequences into 3100 individuals. Case-control status was assigned to individuals in the population based on causal SNVs (cSNVs) randomly sampled from the middle 900-1100 kbp of the 2 Mbp candidate genomic region. For cSNVs, the risk of disease increases according to a logistic-regression model

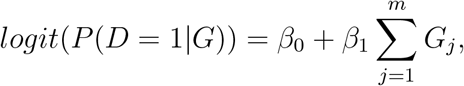

where

- 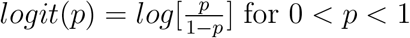 for 0 < *p* < 1,
- D is the disease status (D=1, case; D = 0, control),
- *G* = (*G*_1_, *G*_2_, …, *G_m_*) is the multi-locus genotype of an individual at *m* causal SNVs, where *G_j_* indicates the number of copies of the derived allele at the jth cSNV,
- *β*_0_ is the intercept of the model and controls the sporadic-disease rate, i.e *P*(*D* = 1|*G* = 0), and
- *β*_1_ is the effect parameter which measures the influence of causal variants on the disease.

#### 2.2.1 Simulations under the null hypothesis

We randomly assigned disease status to the 3100 individuals in the population. To ensure a disease prevalence of 5% in the population, 155 out of 3100 individuals were randomly assigned as disease-affected individuals. To form our case-control sample, we randomly sampled 50 cases from the 155 affected individuals and 50 controls from the 2945 unaffected individuals in the population.

#### 2.2.2 Simulations under the alternative hypothesis

We used the disease-trait model above with parameter values set to ensure a high penetrance and low phenocopy rate consistent with genetic-linkage studies (Ott et al., 2015). In particular, we set *β*_0_ = −10 so that the phenocopy rate 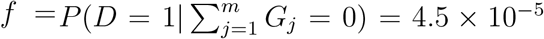, and *β*_1_ = 16 so that the genetic penetrance 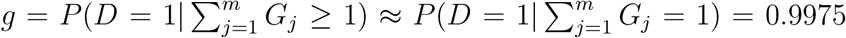. The penetrance ratio was therefore *g/f* ≈ .9975/4.5 × 10^−5^ = 22167.

We aimed for an allelically heterogeneous disease with 15 rare cSNVs of roughly equal frequency in the population. To achieve the targeted disease prevalence, each cSNV had a population allele frequency around 0.16 percent (about 10 copies in the population of 6200 sequences). When necessary, additional very rare variants were chosen to be causal to attain the targeted 5% disease prevalence. Further details about the selection procedure for causal variants can be found in Appendix A.2. After assigning disease status to the 3100 individuals in the population, we randomly sampled 50 cases from the affected individuals and 50 controls from the unaffected individuals. We then extracted the SNV sequences of the case-control sample for analysis.

### 2.3 Genotypic association

We consider Fisher’s exact test and an optimized sequence-kernel association test (Lee et al., 2012). These methods test for association between the trait and the genotypes, either one-at-a-time or in aggregate, and do not consider the relatedness of sequences.

#### 2.3.1 Fisher’s exact test

We use a standard Fisher’s exact test of disease association with genotype frequencies for each SNV implemented in the stats package in base R. Specifically, each of the SNV sites is tested for an association with the disease outcome using a 2 × 3 table to compare the genotype frequencies. At each SNV, our test statistic is the exact p-value, expressed in the negative, base-10-logarithmic scale.

#### 2.3.2 SKAT-O

Single-variant association tests such as Fisher’s exact test have limited power to detect rare variants (Lee et al., 2014). To improve power, aggregation methods, such as the sequence-kernel association test, collapse variants in a window of SNVs into a one-number summary that is then used to test for association. We consider the adaptive aggregation test SKAT-O (Lee et al., 2012) which finds the optimal linear combination of the burden test (Madsen and Browning, 2009) and the sequence kernel association test in terms of power. We applied the SKAT-O test implemented in the SKAT R package (Wu et al., 2011) to each SNV, using a window size of 21 SNVs (~ 14 — 15 kbp in the simulated datasets). The window includes the target SNV at the center and 10 SNVs to the right and left. Target SNVs at either edge of the candidate region had a smaller window size than 21. For example, the window centered at the first SNV has no SNVs to the left and 10 SNVs to the right and thus contains 11 SNVs in total. At each SNV, we record the *p*-value, expressed in the negative-base-10-log scale, as the test statistic.

### 2.4 Descent-based association

Rather than associating genotype frequencies with trait values, we propose instead a linkage analysis that associates similarity in sequence relatedness with similarity in trait values. Since sequences carrying a causal variant tend to cluster on the gene genealogy around the variant, we expect sequence relatedness and trait similarity to be associated in genomic regions harbouring causal variants. As the true gene genealogy is unknown, we reconstruct sequence partitions on the genealogy from the sequence data and calculate distances on these partitions. We then calculate trait dissimilarities between sequences and use them to assess the association between the clustering of sequences and trait values.

#### 2.4.1 Sequence partitions and their distances

To reconstruct sequence partitions, we apply the clustering methods implemented in the R package, `perfectphyloR` (Karunarathna and Graham, 2019). The package takes the sample sequences and returns a perfect phylogeny for a focal SNV. The perfect phylogeny is a rooted tree that recursively partitions DNA sequences (Gusfield, 1991). These nested partitions provide insight into the relatedness of sequences around a focal SNV. Sequences descending from a common ancestral mutation tend to cluster together in a partition. We use the `perfectphyloR` function `reconstructPPregion()` with a minimum window size of 500 variants to reconstruct partitions across the 2-Mbp genomic region. We use a large window size to help resolve non-identical sequences in the reconstruction. Note that the sequence partitions provide no information on coalescence times or the ordering of non-nested coalescence events and so are not genealogical trees. They do however provide information on the nested structure of sequence clusters and therefore on sequence relationships.

At each SNV, we measure the scaled pairwise distances between sequences on the partitions as described in Burkett et al. (2014). These distances measure how closely sequences are related around a focal SNV. To compute the pairwise distances, we apply the `perfectphyloR` function `rdistMatrix()`. Partitions may change along the genome due to recombination. As a result, the pairwise distances may differ for different focal SNVs. A small example of sequence distances on a partition is presented in Appendix A.3.

Phenotypic distances are computed as described in Beckmann et al. (2005). These distances measure the trait dissimilarity of sequences. Briefly, the phenotypic distance between sequence *i* and *j* is defined to be *d_ij_* = 1 – *S_ij_*, where *s_ij_* = (*y_i_* – *μ*)(*y_j_* – *μ*) is the phenotypic similarity score between sequence *i* and *j, y_i_* is the binary phenotype (0 for control or 1 for case), and *μ* is the disease prevalence in the population. For our disease prevalence of 5%, the phenotypic distances are essentially dichotomous. In one group, the distances between case sequences take on the same low values while, in the other group, the distances between control sequences or between case and control sequences take on similar high values.

#### 2.4.2 Measures of association

We associate sequence and phenotypic distances in two ways: via the distance correlation (Székely et al., 2007) or the Mantel coefficient (Mantel, 1967). The distance correlation measures non-linear dependence between two random vectors but can be expressed in terms of pairwise Euclidean distances (Josse and Holmes, 2016). In contrast, the Mantel coefficient measures linear dependence between elements of two distance matrices which do not necessarily have to be Euclidean. At each SNV, we record the distance correlation or Mantel coefficient as the test statistic.

### 2.5 Scoring detection

Through simulation, we compare the abilities of the two genotypic and two descent-based methods to both *detect* association and *localize* causal SNVs. For detection, we are interested in finding *any* association across the entire candidate region. For localization, we are interested in mapping the locus harboring causal variants. We describe a global test to detect association across the entire region and the empirical distribution function (EDF) to graphically compare the resulting global tests. We also describe how we compute the type-I error rate and power of the global tests.

#### 2.5.1 Global tests

For each dataset, we use the maximum test statistic across all the SNVs to obtain a global test statistic across the candidate genomic region. We obtain the null distribution of this global test statistic by randomly permuting the case-control labels of the individuals 1000 times. The *p*-value for the global test is defined as the proportion of test statistics that are greater than or equal to the observed value. The nominal level of all tests is 5%.

##### 2.5.2 Empirical distribution functions

To compare the distribution of *p*-values for each of the methods, we plot their empirical distribution functions or EDFs. The EDF at any point *x* ∈ (0,1) indicates the proportion of simulated datasets with a p-value less than or equal to *x*. Therefore, any method with higher EDF at *x* has a larger proportion of simulated datasets with *p*-value less than or equal to *x*.

##### 2.5.3 Type-I error rate and power

The estimated type-I error rate and power of each method is respectively the proportion of the 500 datasets simulated under the null or alternative hypothesis that are rejected at level 5%. Type-I error rate and power can be extracted from the EDF. For example, when datasets are simulated under the alternative hypothesis, any method with higher EDF at *x* = 0.05 appears to be more powerful at level 0.05. To assess whether the power of two methods differs, we apply McNemar’s test (McNemar, 1947) to the EDFs evaluated at *x* = .05. We use McNemar’s test to account for dependence in test results from the same dataset.

We are particularly interested in the type-I error rate of the Mantel test because it is known to be biased (i.e to have inflated type-I error rate) when the units being permuted are non-exchangeable under the null hypothesis (Guillot and Rousset, 2013). In our context, the sample sequences are not exchangeable owing to their underlying ancestry. However, under the null hypothesis of no association, the casecontrol status of the individuals being permuted is exchangeable. We use a normal approximation to the binomial distribution to obtain an approximate 95% confidence interval for the type-I error rate.

#### 2.6 Scoring localization

To evaluate the localization ability of each method, we calculate the distance of the maximum absolute association signal from the causal region, in base pairs. If more than one maximum is encountered, we take the average of all maxima. We then calculate the EDF of these average distances for the 500 simulated datasets. The EDF at any point 0 ≤ *x* ≤ 2000 kbp gives the proportion of simulated datasets with peak association signal within x kbp of the causal region. Therefore, any method with higher values of the EDF at a given value *x* appears to localize better, within a distance of *x* kbp. To assess whether the localization ability of two methods differs, we apply McNemar’s test to the EDFs evaluated at *x* = 0.

#### 2.7 Post-hoc labeling of case sequences

We propose a procedure to label case sequences as potential carriers or non-carriers of causal variants. Our approach relies on the concept of a genealogical nearest neighbor or GNN (Kelleher et al., 2019b). GNNs arise from the topological properties of genealogical trees, as summarized by the sequence partitions. Case sequences that carry a given rare variant are descended from a common ancestral mutation that arose relatively recently back in time. Therefore, we expect these case sequences to cluster in the sequence partition as GNNs. The GNN proportion of a sequence for a given partition is the proportion of its nearest neighbors in the partition that are case sequences. We then average this proportion over all sequence partitions along the genomic region to obtain an average GNN proportion. A worked example of the calculation of average GNN proportions is illustrated in Appendix A.4. Briefly, any case sequence whose average GNN proportion is consistent with the distribution of GNN proportions in controls is declared to be more closely related to controls than to cases. We group case sequences into carriers and non-carriers of causal variants according to their average GNN proportion. We consider the median of the distribution of GNN proportions in control sequences as our threshold. Specifically, any case sequence with an average GNN proportion less than the median of the average GNN proportion in control sequences is labelled as a non-carrier. We refer to this grouping as the *GNN labeling* of case sequences.

Since we have simulated the sequences and genealogies, we know the true carrier status of each sequence. Therefore, we can compare the accuracy of our GNN labeling to naive labeling, in which all case sequences are assumed to be carriers. Table 1 presents an example confusion matrix for the carrier status of *N* = *a* + *b* + *c* + *d* case sequences in a simulated dataset. Referring to this confusion matrix, we see that the observed misclassification rate is 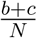.

**Table 1:** Example confusion matrix of carrier status for *N* case sequences

|  |  | GNN-predicted status |  |
| --- | --- | --- | --- |
|  |  | Non-Carrier | Carrier |
| True status | Non-Carrier | a | b |
|  | Carrier | c | d |

$N=a+b+c+d$

## 3 Results

To start, we present an analysis of an example dataset to give insight into the association methods. We then present estimated type-I error rates and rates of *detection* and *localization* of the causal region. Finally, we present misclassification error rates for the proposed post-hoc labeling of case sequences.

### 3.1 Example dataset

We first summarize the causal variants in the population and sample. Next, we show profiles of various association statistics across the candidate genomic region and apply the proposed procedure for post-hoc labeling of the case sequences.

#### 3.1.1 Population and sample summaries

The population of 3100 individuals (6200 sequences) has 4723 SNVs in a 2 Mbp genomic region. Among these SNVs, 2904 are segregating in the sample of 50 case and 50 control individuals, including all 15 cSNVs. In the population, all sequences and all but one individual carry zero or one cSNV. The one individual with two carrier sequences is included in the sample as a case. Table 2 summarizes the causal variants in the sample and population. The column labeled “Position (kbp)” gives the physical position of cSNVs along the genome in kbp. The columns labeled “Population” and “Sample” count the number of case and control sequences that are carrying any causal variants in the population and sample, respectively. The column labeled “DAF” gives the derived allele frequency of causal variants in the population, expressed as a percentage. The causal variants are all rare with a maximum population DAF of 0.19%. A total of 154 and 51 case sequences carry a cSNV in the population and sample, respectively. None of the control sequences carry cSNVs.

**Table 2:** Summaries of causal variants in the sample and population.

| cSNV | Position<br>(kbp) | Population sequences |  |  | Sample sequences |  |
| --- | --- | --- | --- | --- | --- | --- |
|  |  | Case<br>(154) | Control<br>(0) | DAF% | Case<br>(100) | Control<br>(100) |
| 1 | 928.761 | 9 | 0 | 0.14 | 4 | 0 |
| 2 | 937.392 | 12 | 0 | 0.19 | 7 | 0 |
| 3 | 940.023 | 12 | 0 | 0.19 | 4 | 0 |
| 4 | 942.571 | 9 | 0 | 0.14 | 5 | 0 |
| 5 | 946.127 | 10 | 0 | 0.16 | 1 | 0 |
| 6 | 993.008 | 12 | 0 | 0.19 | 4 | 0 |
| 7 | 994.439 | 10 | 0 | 0.16 | 3 | 0 |
| 8 | 998.710 | 11 | 0 | 0.18 | 4 | 0 |
| 9 | 1002.568 | 9 | 0 | 0.14 | 2 | 0 |
| 10 | 1003.525 | 11 | 0 | 0.18 | 2 | 0 |
| 11 | 1016.514 | 9 | 0 | 0.14 | 1 | 0 |
| 12 | 1039.256 | 10 | 0 | 0.16 | 4 | 0 |
| 13 | 1045.524 | 11 | 0 | 0.18 | 3 | 0 |
| 14 | 1054.265 | 10 | 0 | 0.16 | 4 | 0 |
| 15 | 1082.301 | 9 | 0 | 0.14 | 3 | 0 |

#### 3.1.2 Association profiles

The association profile is a scatter plot with genomic coordinates on the horizontal axis and SNV-specific measures of association on the vertical axis. Figure 1 presents the association profiles of different methods for the example dataset.

**Figure 1:**
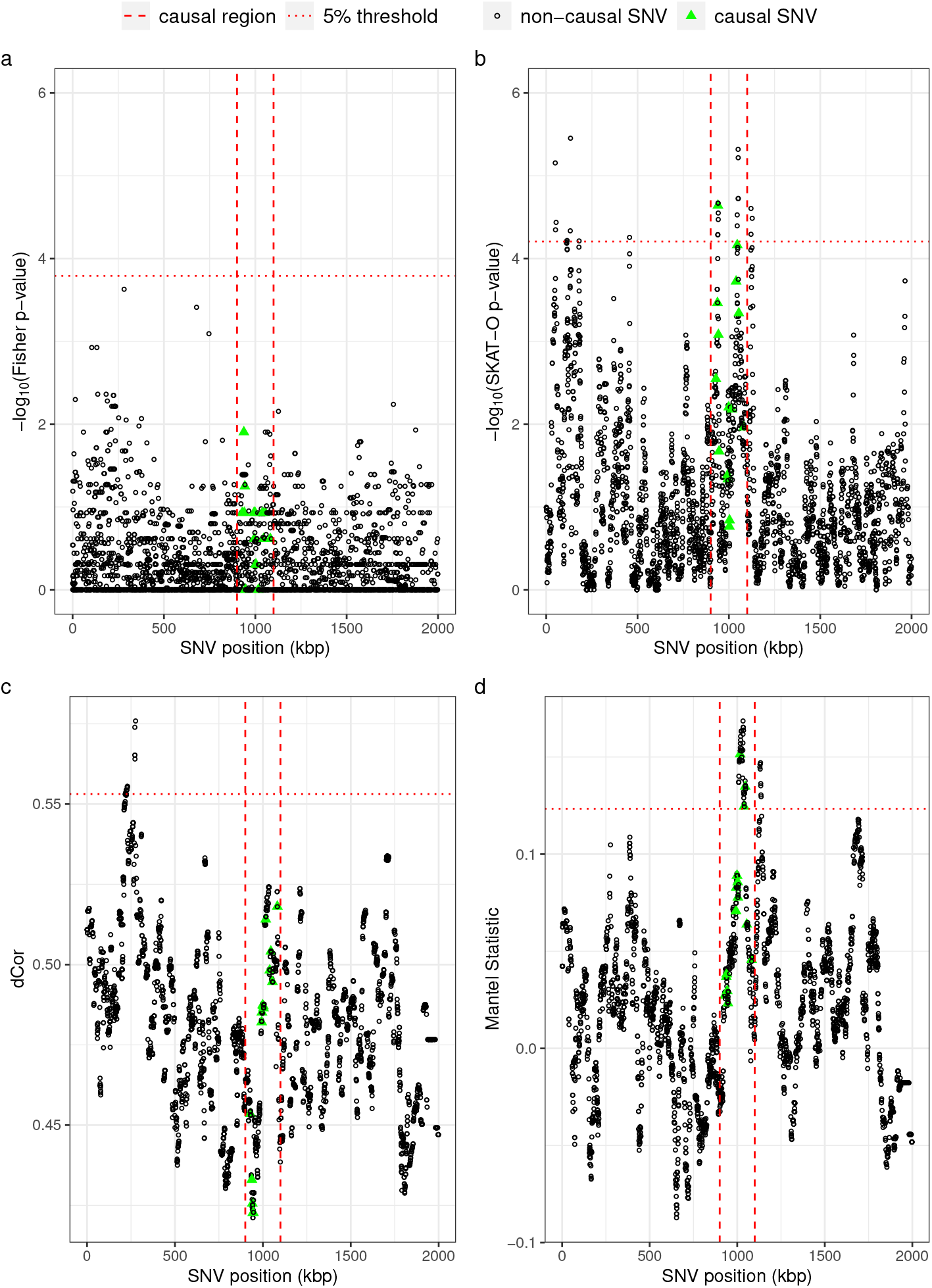
Association profiles, a) Fisher’s exact test (FET), b) SKAT-O, c) distance correlation (dCor), and d) Mantel. The vertical red-dashed lines indicate the region from which the causal SNVs were selected. The green triangles represent the causal SNVs. The maximum value of SNV-specific statistics over the entire genomic region is used in a permutation test for the presence of any association. The horizontal dotted line represents the 5% significant threshold based on 1000 permutations of the individual disease phenotypes. The detection *p*-values for FET, SKAT-O, dCor, and Mantel are 0.089, 0.005, 0.008, and 0.002 respectively.

The *x*-axes in all panels is the genomic position in kbp. The *y*-axes show either a transformed SNV-specific *p*-value (genotypic association methods), or an SNV-specific measure of association (descent-based association methods). The vertical red-dashed and horizontal red-dotted lines indicate, respectively, the causal region from which the cSNVs were randomly selected and the 5% significant threshold for the global test of any association. The Mantel, SKAT-O and distance correlation tests detect significant association, but the Mantel test is the only method that correctly localizes the causal region. The profiles for Fisher’s exact test and distance correlation in panels (a) and (c) appear similar, and the peak association signal of both methods occurs in approximately in the same genomic position. We will return to this point later when discussing the simulation results.

#### 3.1.3 Post-hoc labeling of case sequences

We apply the proposed GNN-labeling procedure to classify case sequences in the example dataset. Figure 2 shows the boxplots of average GNN proportions of sequences grouped by their status as case carriers or case non-carriers of causal variants and controls. We use the median of average GNN proportion in control sequences to classify the case sequences into carriers and non-carriers. In the example dataset, all but three of the true carriers are correctly predicted by the GNN labeling.

**Figure 2:**
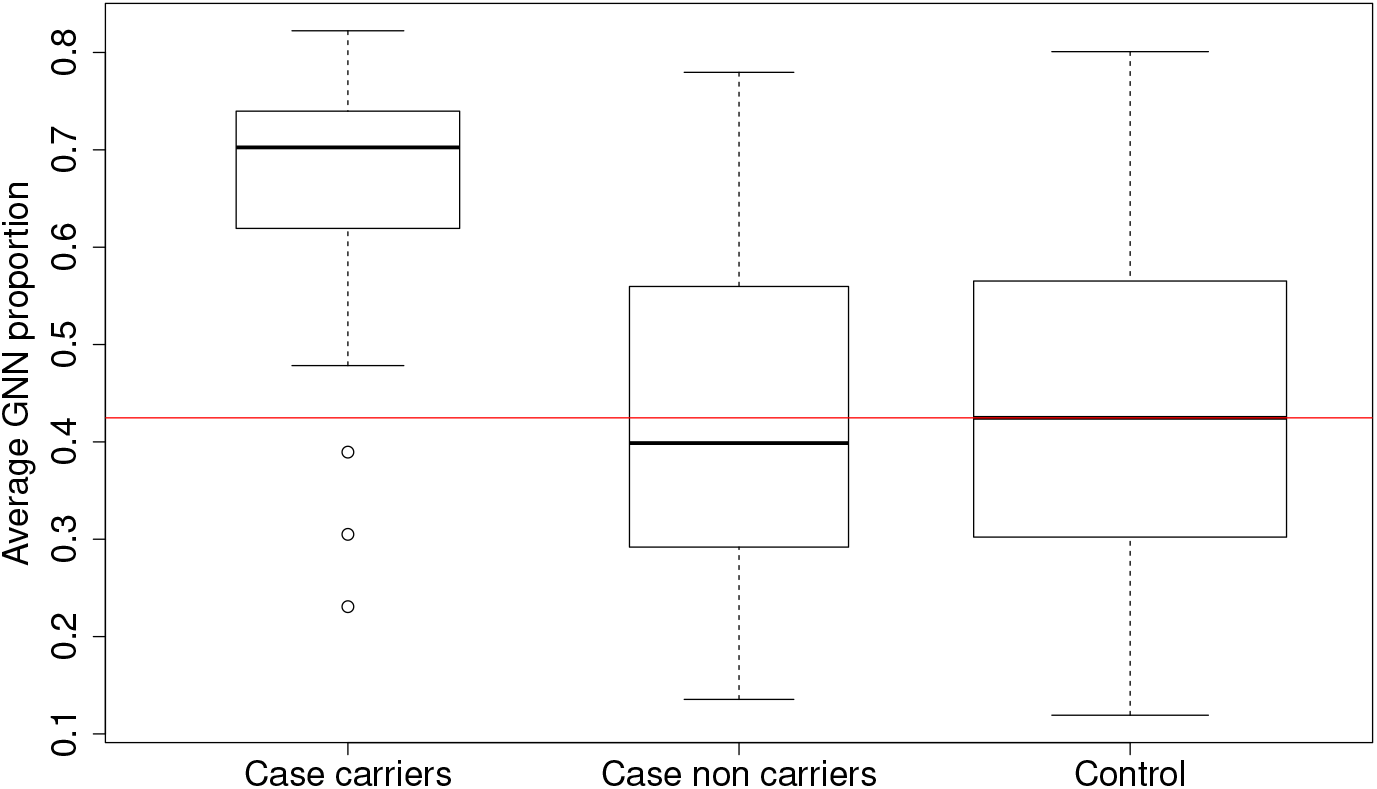
Average GNN proportions of sequences grouped by their status as case carriers of causal variants, case non-carriers of causal variants or controls. The horizontal red line is the median of the average GNN proportion in control sequences.

Tables 3 (a) and (b) show the confusion matrices for naive and GNN labeling, respectively. Naive labeling considers all 100 case sequences to be carriers of a cSNV. The observed misclassification rates for naive and GNN labeling are 49% and 25%, respectively. Post-hoc, GNN labeling therefore improves the identification of carriers of a cSNV among case sequences.

**Table 3:** Carrier status for *N* = 100 case sequences in the example dataset using a) naive labeling and b) GNN labeling.

(a) Naive labelling
|  |  | Naive status |  |
| --- | --- | --- | --- |
|  |  | Non-Carrier | Carrier |
| True status | Non-Carrier | 0 | 49 |
|  | Carrier | 0 | 51 |

N=100 (b) GNN labelling
|  |  | GNN-predicted status |  |
| --- | --- | --- | --- |
|  |  | Non-Carrier | Carrier |
| True status | Non-Carrier | 27 | 22 |
|  | Carrier | 3 | 50 |

N=100

Table 4 considers the 51 case sequences that carry a cSNV in the example dataset, and presents the number that are correctly predicted by GNN labeling. Three carrier case sequences are incorrectly predicted, corresponding to the cSNVs in highlighted rows of the table. As the case sequences carrying cSNVs 5 and 11 are sample singletons, we would not expect their genealogical nearest neighbours to be overrepresented by case sequences. Thus, we would not expect the GNN labeling to correctly predict their carrier status.

**Table 4:** Carrier case sequences for each cSNV and number predicted by GNN labeling.

| cSNV | Sample sequences |  |
| --- | --- | --- |
|  | Case | GNN predicted |
| 1 | 4 | 4 |
| 2 | 7 | 7 |
| 3 | 4 | 4 |
| 4 | 5 | 5 |
| 5 | 1 | 0 |
| 6 | 4 | 4 |
| 7 | 3 | 3 |
| 8 | 4 | 4 |
| 9 | 2 | 2 |
| 10 | 2 | 2 |
| 11 | 1 | 0 |
| 12 | 4 | 4 |
| 13 | 3 | 3 |
| 14 | 4 | 3 |
| 15 | 3 | 3 |

### 3.2 Detection

We present the simulation results for type-I error rates first, then the results for power.

#### 3.2.1 Type-I error rate

To estimate type-I error rates, we considered 500 datasets simulated under the null hypothesis of no association with cSNVs. Figure 3 shows the empirical distribution functions (EDFs) of the permutation *p*-values from a global test of association across the entire genomic region, for each of the association methods. In both panels, *p*-values are labeled in the natural scale but plotted in the log-10-scale. The *y*-axis in the left panel of the figure is shown up to 0.30. The right panel of the figure magnifies EDFs around the 5% significance level. Figure 4 presents point and approximate 95%-confidence interval estimates of the type-I error rates. The results of both figures suggest that the type-I error rates of all association methods are controlled at the nominal 5% level. Numerical values for the point and 95% confidence-interval estimates of the type-I error rates are reported in Table A.1 in Appendix A.5.

**Figure 3:**
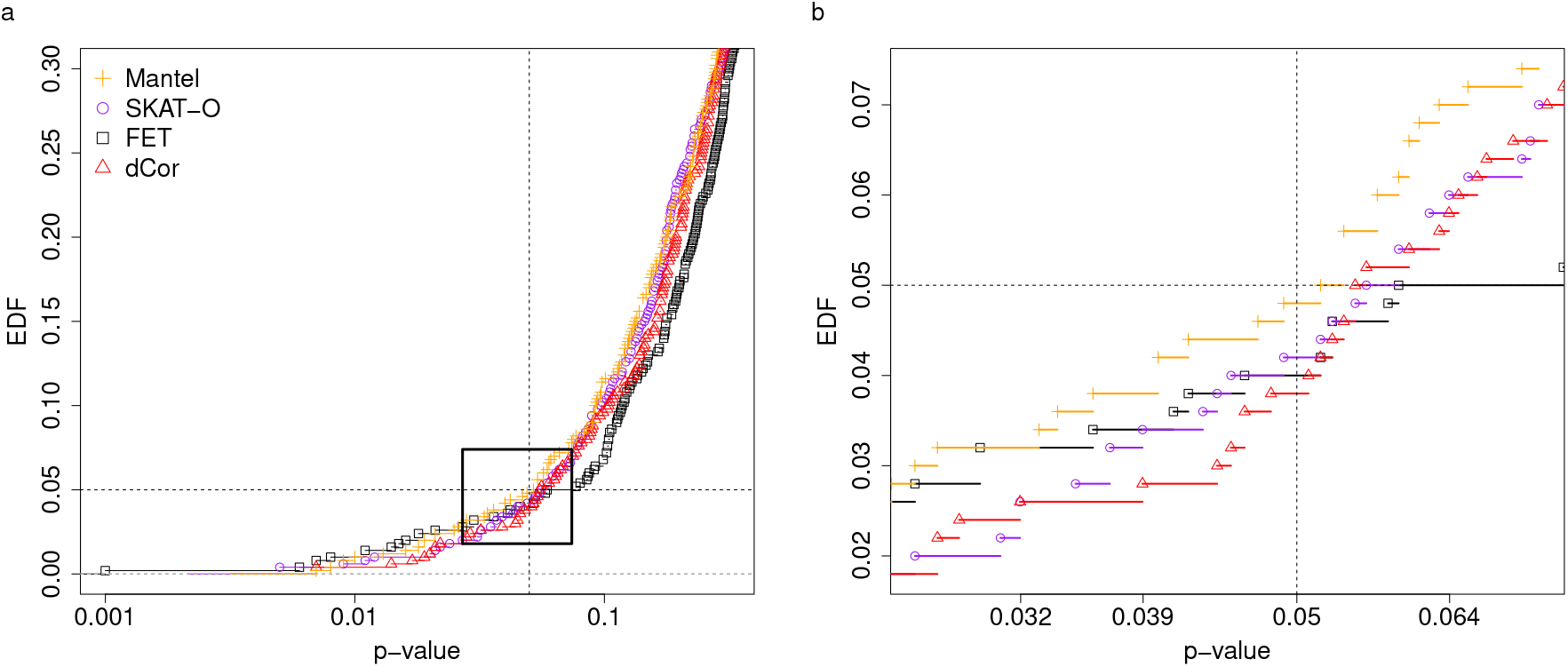
Empirical distribution functions (EDFs) of permutation *p*-values from a global test of association across the genomic region. Four methods are compared: Fisher’s exact test (FET), SKAT-O, distance correlation (dCor) and Mantel. a) Original b) Zoomed version. On the *x*-axis, *p*-values are labeled in the natural scale but plotted in the log-10-scale. The vertical and horizontal dashed lines indicate the nominal 5% level.

**Figure 4:**
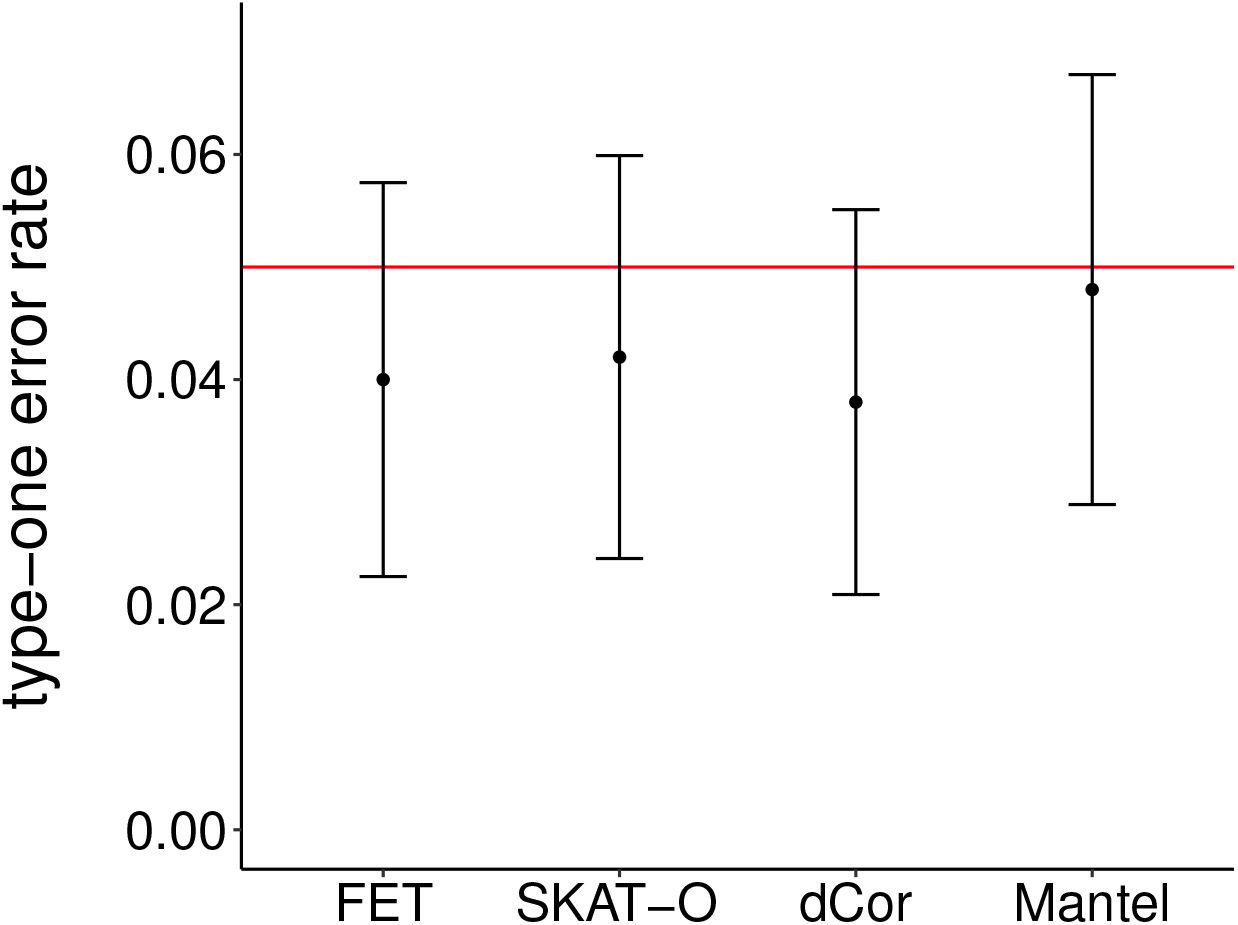
Point and approximate 95%-confidence interval estimates for type-I error rate in Fisher’s exact (FET), SKAT-O, distance correlation (dCor), and Mantel tests. The horizontal dashed line is the nominal 5% level.

#### 3.2.2 Power

The EDFs shown in Figure 5 are based on 500 datasets that have been simulated under the alternative hypothesis of association with causal SNVs. The EDFs suggest that the SKAT-O and Mantel tests outperform the other tests for detecting the association signal. The performance of the SKAT-O and Mantel tests at level 5% does not differ significantly (McNemar *p*-value = 0.60).

**Figure 5:**
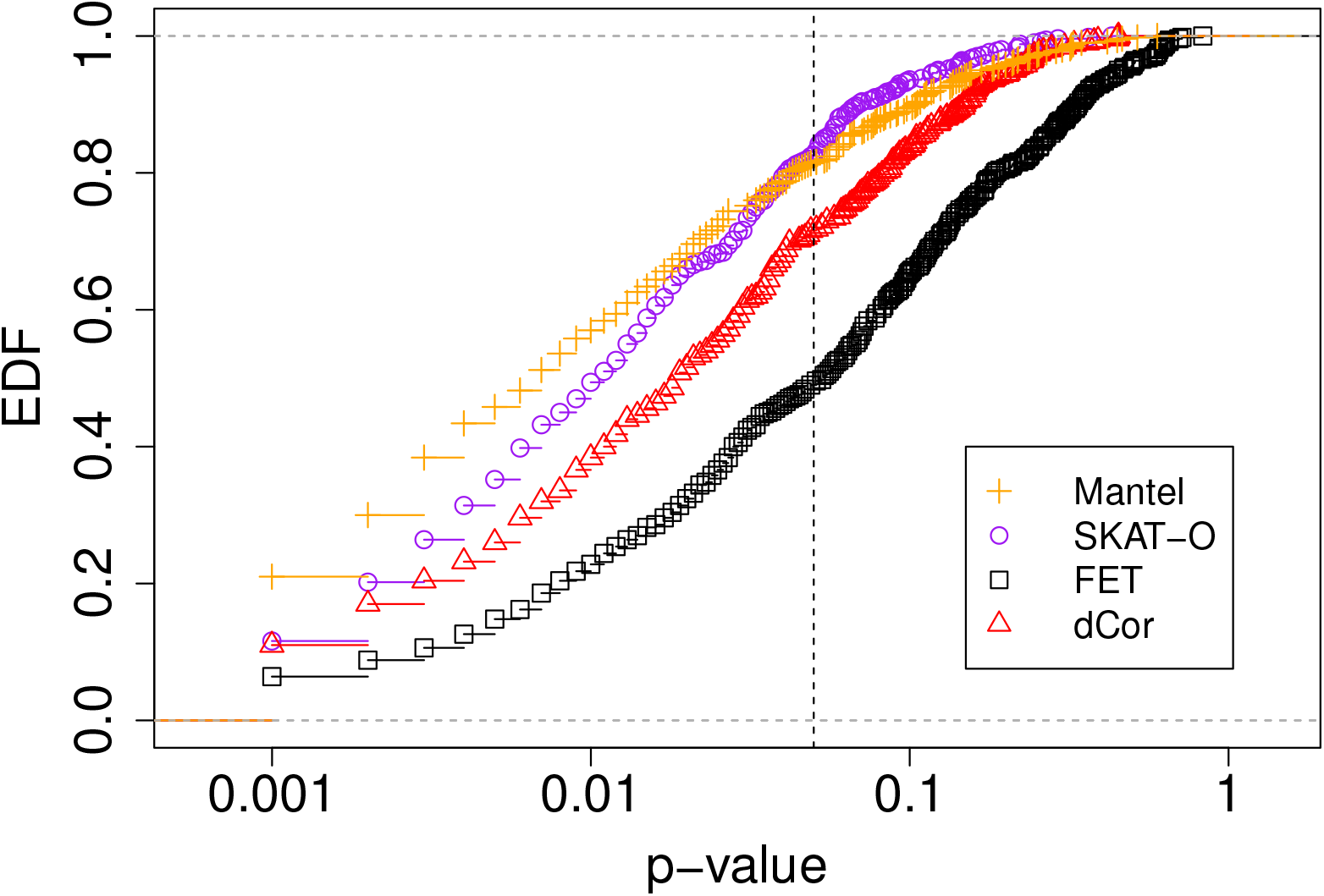
The empirical distribution function (EDF) plot of permutation *p*-values from a global test of association across the genomic region. Four methods are compared: Fisher’s exact test (FET), SKAT-O, distance correlation (dCor), and the Mantel statistic. On the *x*-axis, *p*-values are labeled in the natural scale but plotted in the log-10 scale. The vertical dashed line indicates a *p*-value of 0.05.

A scatter plot of detection *p*-values from the SKAT-O and Mantel tests is shown in Figure 6. The Pearson correlation between these two sets of *p*-values is 0.175 and differs significantly from zero (*p* < 0.0001). In 62 of the 500 datasets, the Mantel test detects the association signal but the SKAT-O test does not (fourth quadrant). In 68 datasets, the SKAT-O test detects the association signal but the Mantel test does not (second quadrant). Both methods detect the association signal in 345 datasets (third quadrant) and neither method detects the signal in 25 datasets (first quadrant). The observed discordance rate between the tests is 130/500, or 26%. These results suggest that the SKAT-O and Mantel tests are picking up on different aspects of the association signal.

**Figure 6:**
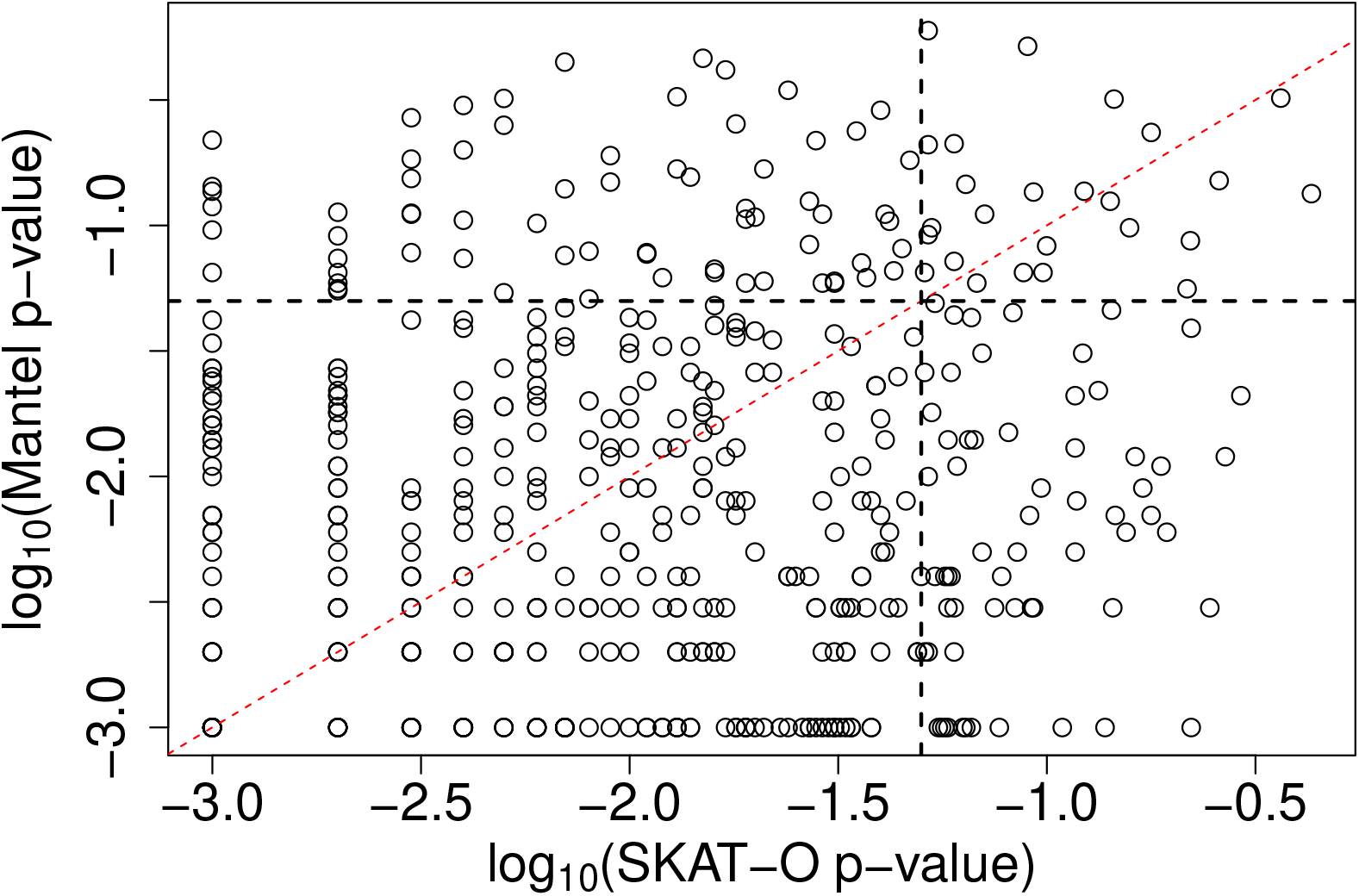
The relationship between *p*-values from the Mantel test and SKAT-O in the log-10 scale. The vertical and horizontal black-dashed lines show *p*-values of 0.05. The Pearson correlation between the transformed *p*-values is 0.175 (p <0.0001). The red-dashed line is *y=x.*

### 3.3 Localization

The EDFs of average distances from the causal region are shown in Figure 7, for the four association methods. The Mantel profile appears to localize the causal region far better than any of the others, followed by SKAT-O. In fact, the Mantel profile localizes significantly better than SKAT-O (McNemar *p*-value = 0.0042)

**Figure 7:**
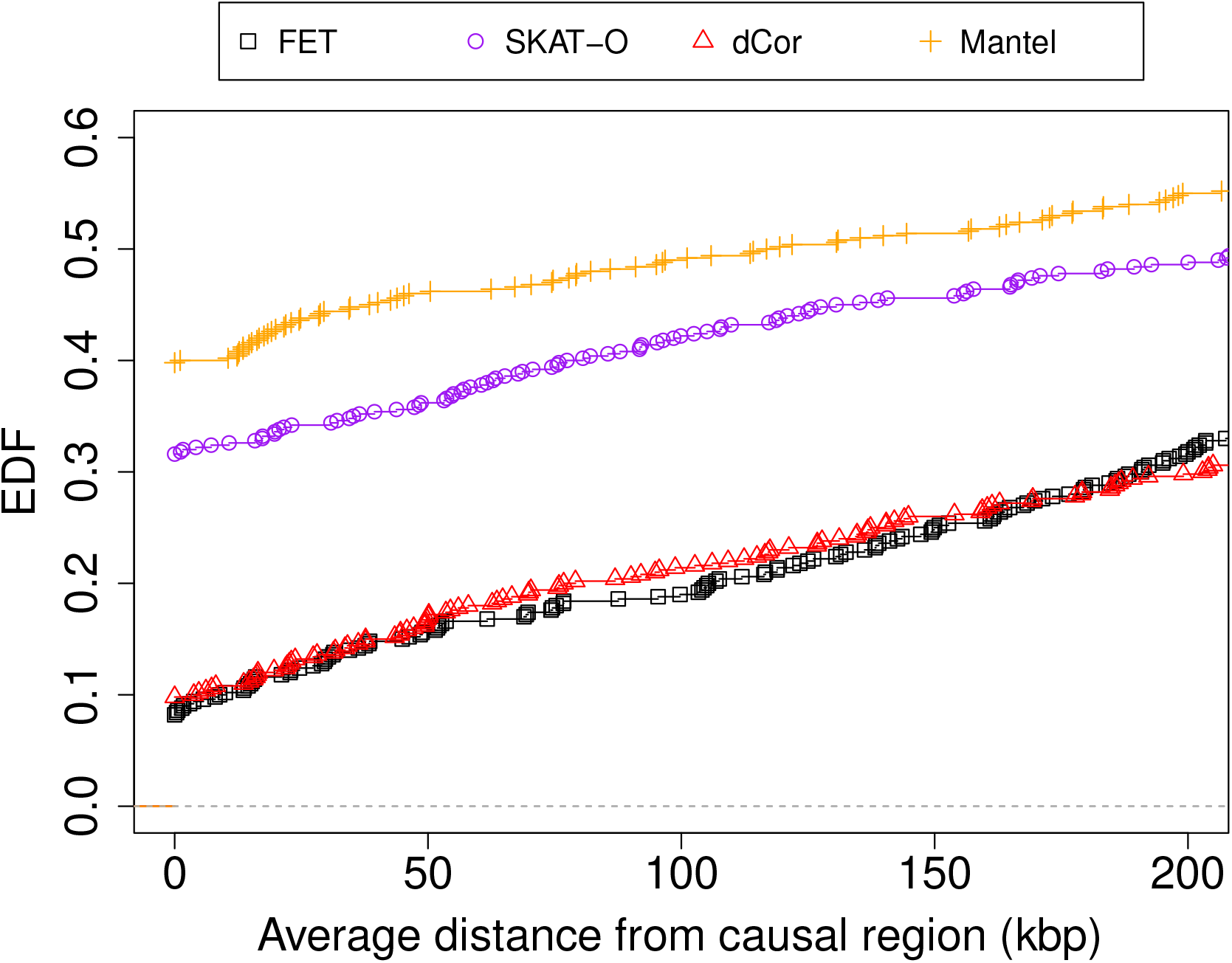
The empirical distribution functions (EDFs) for the average distance of the peak association signal from the causal region, for 500 datasets simulated under the alternative hypothesis of association. Four methods are compared: Fisher’s exact test (FET), SKAT-O, distance correlation (dCor) and Mantel. To make the comparison easier and for better resolution, the *x*-axis is shown only for genomic distances less than 200 kbp.

From the figure, we see that Fisher’s exact test and the distance correlation localize about 10% of the 500 simulated datasets to the causal region. However, the causal region comprises 10% of the candidate region being fine-mapped, and so these two methods are localizing no better than random chance. In the example dataset, Fisher’s exact test and the distance correlation had similar association profiles which localized the peak signal to roughly the same genomic position. To investigate the co-localization properties of the methods, we calculated the Pearson correlation of their average distances from the causal region. Amongst all pairs of methods, the maximum correlation of 0.30 (p-value ≈ 0) belongs to Fisher’s exact test and the distance correlation. Our findings suggest that Fisher’s exact test and distance correlation tend to co-localize the association signal more than any other pair of methods considered.

### 3.4 Performance of case-sequence labeling

We use the 500 datasets simulated under the alternative hypothesis to compare the performance of GNN labeling of case sequences to a naive labeling scheme in which all case sequences are assumed to be carriers of cSNV. For the 100 sampled case sequences in each dataset, we compute the misclassification rates of the labeling procedures. Figure 8 shows the scatter plot of these rates for the 500 datasets, with the GNN rates on the vertical axis and the naive rates on the horizontal axis. The naive misclassification rates on the horizontal axis have only four values (0.48, 0.49, 0.50, and 0.51), which have been randomly perturbed for better viewing. From the red-dashed line indicating *y* = *x*, we can see that GNN labeling has a uniformly lower misclassification error rate than naive labeling, across all 500 datasets. Thus, post-hoc labeling of case sequences by the GNN procedure may be of practical use for predicting which case sequences carry causal variants.

**Figure 8:**
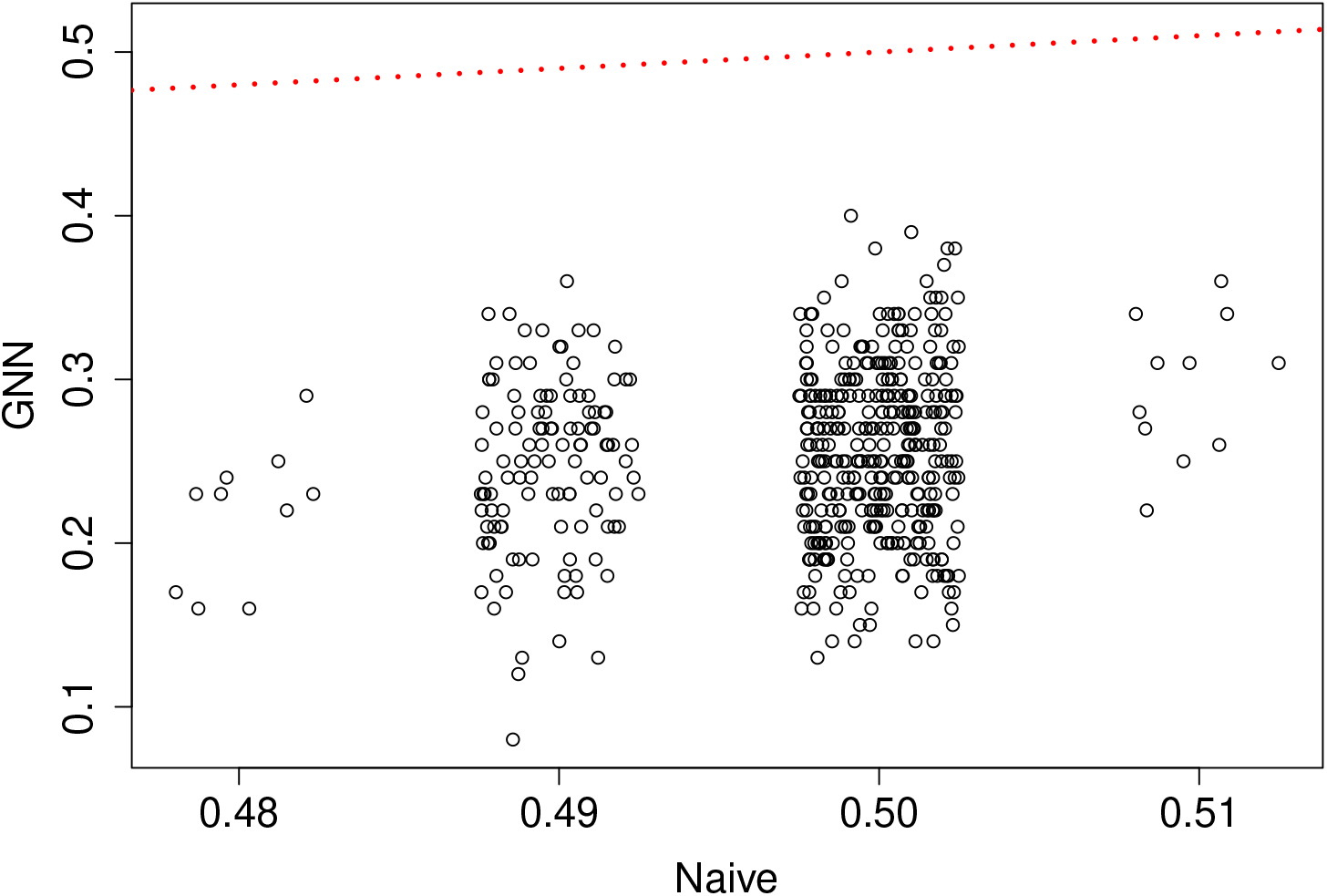
Misclassification error rate of cSNV carrier status in case sequences, for GNN versus Naive labeling across 500 simulated datasets. The red-dashed line is *y = x*.

## 4 Discussion and Conclusion

We have explored the feasibility of linkage fine-mapping on sequences for an allelically heterogeneous disease. In particular, we have compared the ability of linkage and genotypic-association approaches to detect and fine-map a causal locus in simulated datasets. The linkage methods we have considered use sequences rather than individuals as the unit of observation. These methods associate similarity in relatedness of sequences with similarity in trait values, using either the distance correlation or Mantel coefficient as a measure of association. For comparison, we include two genotypic-association methods, a single-variant Fisher’s exact test and the SKAT-O test that aggregates variants. While our linkage methods use haploid sequences and the genotypic association methods use diploid individuals as the unit of observation, *both* classes of methods assume a diploid disease model. A consequence for the linkage methods is that some case individuals will have sequences that do not carry causal variants. We therefore introduce a *post-hoc* labeling procedure to group case sequences into carriers and non-carriers of causal variants, inspired by the idea of genealogical nearest neighbors (GNN) described in Kelleher et al. (2019b). We view this simulation investigation as a proof of principle illustrating the potential of sequence-based linkage approaches. A future direction of research would be to compare the linkage and SKAT-O approaches on real datasets from fine-mapping studies.

Our example data analysis and simulation study indicate that sequence-based linkage methods are useful for population-based fine-mapping of an allelically heterogeneous disease. In particular, we find that: (i) a linkage-based Mantel test detects rare causal variants as well as a state-of-the-art genotypic-association test, SKAT-O; (ii) the Mantel association profiles best localize rare causal variants among all the methods considered; and (iii) GNN labeling of case sequences is helpful for removing sequences that do not carry causal variants. These findings suggest the following strategy for fine-mapping an allelically heterogeneous disease in case-control samples. First, detect disease association with either the Mantel or SKAT-O test. Once a disease association has been detected, localize the causal region with the Mantel association profile and refine the case sequences by removing those labeled as noncarriers by the GNN procedure. The putative causal locus and putative carrier case sequences can then be searched for causal variants. Our work extends earlier investi-gations of sequence-based linkage mapping that relied on the known gene genealogies (Burkett et al., 2014; Karunarathna and Graham, 2018). Instead, we infer the topological structure of unknown genealogies from sequence data. Our sequence-based approaches are therefore practical for linkage fine-mapping with population-based data.

We evaluated the type-I error rate using datasets simulated under the null hypothesis of no association. Our results suggest that the type-I error rate of all the methods is well controlled at the 5% nominal level. The Mantel test was of particular interest because it has been criticized for inflated type-I error rates when the units being permuted are non-exchangeable (Guillot and Rousset, 2013). In our context, however, the size of the Mantel test is maintained because the case and control status of individuals is exchangeable under the null hypothesis. Fisher’s exact test had the lowest power of all the methods, as expected for a single-variant method detecting rare variants. The Mantel test of linkage and the SKAT-O test of genotypic-association had the highest power of all methods. The estimated power of these tests at level 5% did not differ significantly (McNemar *p*-value = 0.60). The similar power of the Mantel and SKAT-O tests prompted us to look into the agreement of their *p*-values across the simulated datasets (Figure 6). The random pattern in the figure as well as the observed discordance rate of 26% between the two significance tests at level 5% suggests that the Mantel and SKAT-O tests detect different aspects of the association signal. The complementary nature of the tests indicates that a combined test, e.g., using Fisher’s method of combining p-values (e.g. Derkach et al., 2013), could be more powerful than either the Mantel or SKAT-O tests alone. Investigating the power of combined tests is an area for future work. The Mantel test had higher detection power than the distance-correlation test (Figure 5). We note that the Mantel test is well suited to a disease of low prevalence (5%) because the trait distances, between case/case pairs on one hand and case/non-case and non-case/non-case pairs on the other, are essentially binary (results not shown). The relationship between trait distances is then essentially a straight line and therefore well captured by the Pearson correlation coefficient. By contrast, the distance correlation assumes Euclidean distances (Josse and Holmes, 2016) and so may be unsuitable for our partition distances between sequences.

The principal finding of our simulation study is that, when the penetrance model favours linkage analysis, the Mantel association profile localizes the causal region *significantly better* than the next-best SKAT-O (McNemar *p*-value = 0.0042). In contrast, Fisher’s exact test and the distance correlation localized the causal region the worst, and in fact did no better than random guessing of the location. Interestingly, among all pairs of methods considered, the average distance of the peak association signal from the causal region was the most highly correlated for the Fisher’s exact and distance-correlation methods (*r* = 0.30, *p*-value≈ 0). In some datasets, the peak association signals for Fisher’s exact test arise from synthetic associations with common SNVs outside the causal region that happen to tag multiple causal SNVs by chance (Dickson et al., 2010). It would be interesting to investigate whether distance correlation is also vulnerable to synthetic associations, given its tendency to co-localize the association signal with Fisher’s exact test.

Throughout, we have applied SKAT-O with a window size of 21 SNVs. This window size corresponds to a genomic region of roughly 14-15 kb, about the size of a typical human gene, in our simulated datasets. We also investigated the detection and localization properties of SKAT-O using windows of 11, 41, 61, and 81 SNVs (results not shown). We found that a SKAT-O window size of 61 SNVs yielded slightly greater detection power than the Mantel test. SKAT-O localization rates improved slightly, up to a window size of 61 SNVs, before falling off for larger window sizes. However, no SKAT-O window size achieved better localization than the Mantel method. Such tuning of the SKAT-O window size is not possible in practice, as it will depend on unknowns such as the presence and location of any causal variants. In addition, SKAT-O already involves optimizing over a linear combination of its constituent burden and variance-component tests, so that optimizing over window size would add extra computational burden. Development of practical and computationally feasible procedures for tuning the SKAT-O window size would be an interesting avenue for further investigation.

Our results suggest that sequence-based linkage analysis is useful for fine-mapping allelically heterogeneous traits. To start the discussion, we have used simulation to explore disease traits in a case-control study design, under a genetic architecture that favors linkage methods. In particular, we simulated high-penetrance, low-frequency causal variants. Examples of diseases influenced by high-penetrance, low-frequency variants are familial breast and ovarian cancer (Bailey-Wilson and Wilson, 2011), familial bipolar disorder (Francks et al., 2010), hearing impairment, familial goitres and familial hypertension (Ott et al., 2015). In the absence of allelic heterogeneity, we do not expect sequence-based linkage methods to offer advantages over genotypic association methods such as SKAT-O, for either detection or localization. By analogy to family-based linkage analysis, lower penetrance ratios are expected to reduce the effectiveness of sequence-based linkage analysis in relation to association approaches such as SKAT-O. Further simulations under a larger variety of allele frequency and penetrance parameters are an important direction for future work.

A related area of future work is to expand the scope to other study designs and genetic architectures. For example, how is sequence-based fine-mapping affected as we vary the type of trait (i.e. binary disease versus quantitative), the type of populationbased study design (e.g. case-control, cohort or cross-sectional sampling), and the number and frequency of causal variants within the trait locus? We would also like to investigate the impact of sequencing errors on the sequence-based linkage methods. Our initial thoughts are that sequencing errors will attenuate the association signal by misclassifying carriers of causal variants as non-carriers or *vice versa*, but is unlikely to create false-positive associations.

Adjusting for confounding variables is another area of interest. One option to deal with confounding is a partial Mantel test of association between two distance matrices given a third (Smouse et al., 1986). This extension would allow testing for association between partition distances and phenotype distances given a third distance matrix based on the confounding variables. In fine-mapping, fine-scale population structure is a confounding variable of particular concern because rare variants tend to cluster geographically due to their recent origin (Mathieson and McVean, 2012). Adjusting for fine-scale population structure when fine-mapping rare variants is challenging, though recently proposed permutation approaches offer a potential way forward (Bouaziz et al., 2021).

Our investigation of sequence-based linkage methods has focused on fine-mapping in a candidate region, but these methods also have the potential to scale up to genome-wide analysis as long as computational resources are available. For example, on a 2.1Ghz Intel processor, and with the sequence partition distances in hand, calculation of the Mantel association profile for the example dataset took 2.18 seconds, and calculation of 1000 further scans for the permutation distribution took about 18 minutes. Scaling from a 2MB region to the entire genome is expected to take about 54.5 minutes for a single scan, and about 27,000 minutes for the permutation replicates. However, permutations are easily parallelized across nodes of a compute cluster. Our study was focused on fine mapping, and so used the `perfectphyloR` R package for partition reconstruction. As `perfectphyloR’s` reconstruction does not scale to genome-wide data, we recommend alternative genome-wide reconstructors such as those implemented in `tsinfer` (Kelleher et al., 2019a) or `Relate` (Speidel et al., 2021). For high-penetrance diseases influenced by multiple low-frequency variants, we expect sequence-based linkage analysis to have similar power to SKAT-O and better localization at the genome-wide scale, as in the fine-mapping results from the current investigation.

## Acknowledgements

C.K. would like to thank Professors Kelly Burkett and Marie-Hélène Roy-Gagnon for helpful comments. We acknowledge the support of the Natural Sciences and Engineering Research Council of Canada (NSERC), RGPIN/04296-2018

## Data Availability Statement

The data supporting the findings of this study are available at DOI: 10.5281/zenodo.6592319. The code used to generate and analyze the data is available at https://github.com/SFUStatgen/PBJ0.

## A Appendices

### A.1 Variant allele frequency spectrum of simulated data

**Figure A.1:**
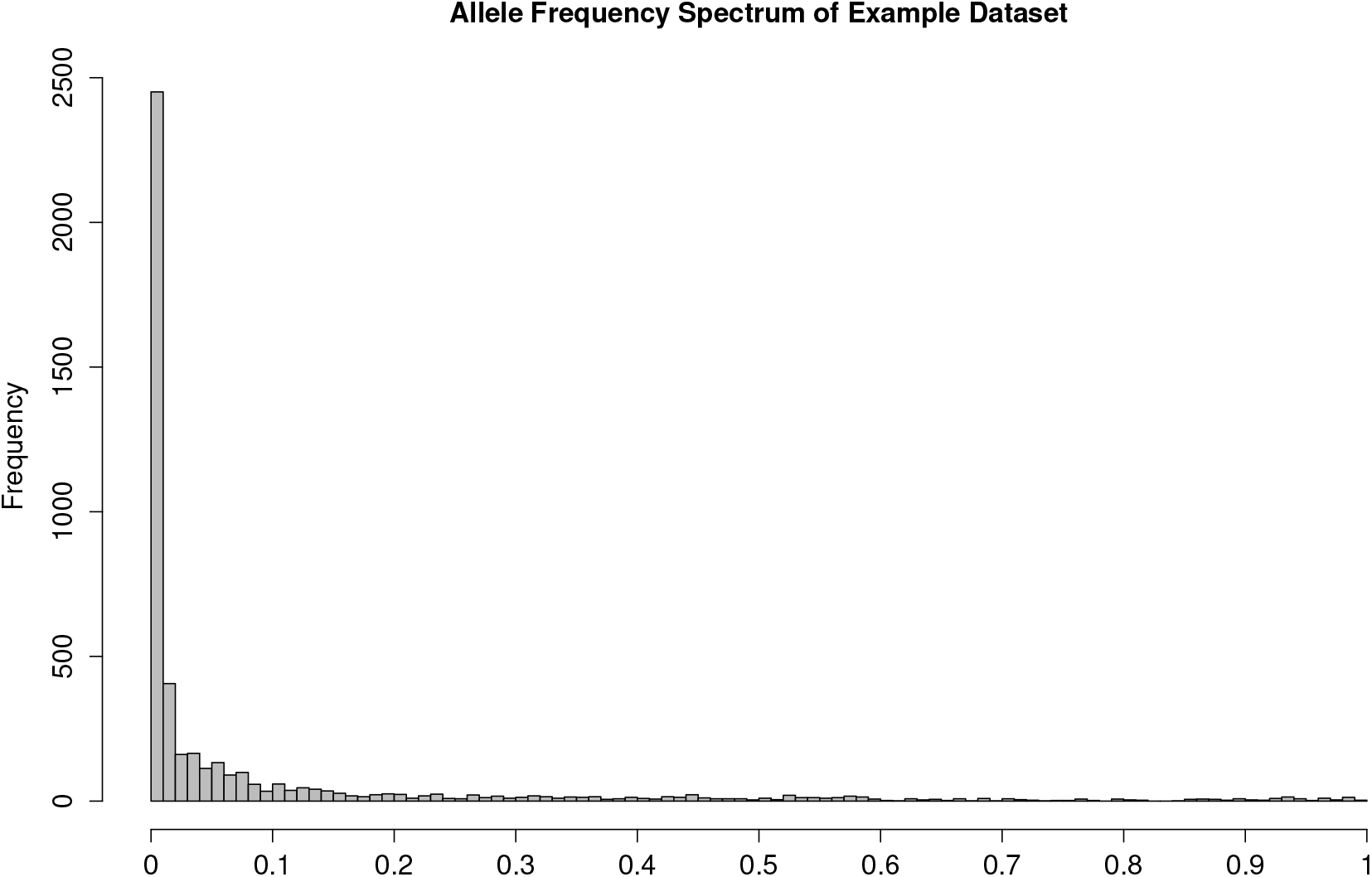
Variant allele frequency spectrum of the example dataset.

### A.2 Causal variant selection

The disease-trait model specifies that 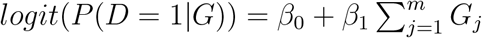 so that

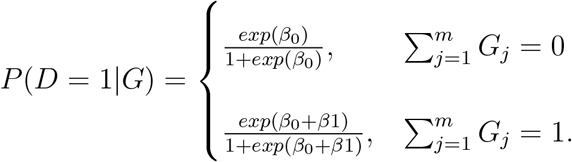

As the cSNVs are rare, we make the simplifying assumption that each individual carries at most one copy of a cSNV. Then the population prevalence of the disease is

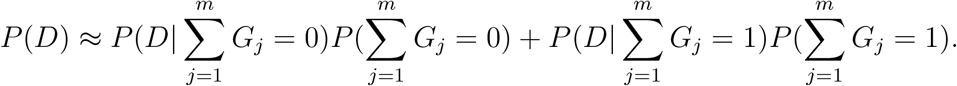

Setting the prevalence to 0.05, we obtain

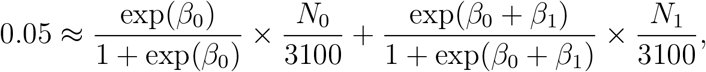

where *N*_0_ and *N*_1_ are respectively the number of individuals in the population that carry zero and one copy of a cSNV. Setting *β*_0_ = −10, *β*_1_ = 16 and *N*_1_ = 3100 — *N*_0_, we obtain *N*_0_ ≈ 2945 and *N*_1_ ≈ 155. We select 15 variants of roughly equal frequency in the population such that their total number of copies is around 155. Thus each cSNV has a frequency of about 155/15 = 10.33 in the population of 6200 sequences.

### A.3 Sequence distances on partition

Figure A.2 shows an example partition with 4 sequences labeled as 1 to 4 and distances assigned from the *rdistMatrix()* function in the *perfectphyloR* package. As illustrated in the figure, the distance between a sequence and its ancestral node is one and the distance between two neighboring nodes that descend from the same most-recent common ancestral node is two.

**Figure A.2:**
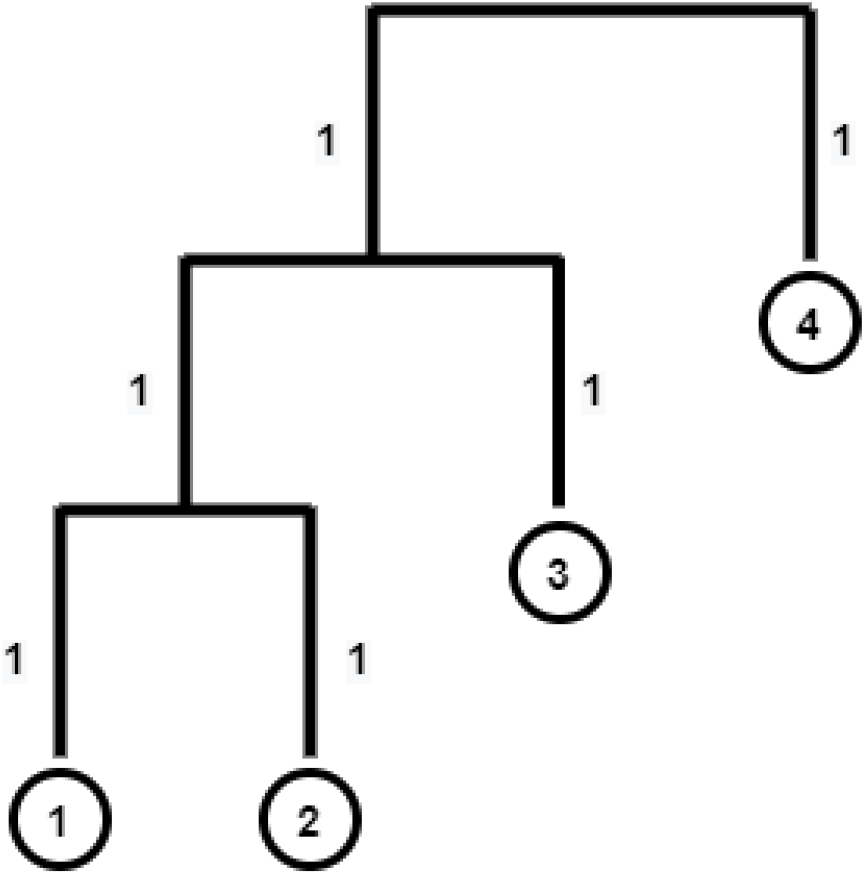
Distance between sequences assigned from the *rdistMatrix()* function in the *perfectphyloR* package. 1, 2, 3, and 4 are sequences.

### A.4 A worked example for calculation of GNN

To illustrate the GNN calculation, we consider a worked example of four sequences of length 10 kbp, as shown in Figure A.3. In the figure, the subregion from 0 kbp to 6 kbp is spanned by partition A, and the rest of the region by partition B. In other words, when we reconstruct the partition at each SNV position within the first 6 kbp, we have only one partition structure (partition A), and the rest of the region has the structure of partition B. The GNN proportion for a given target sequence can be computed as follows. For example, suppose we choose sequence 1 as our target sequence. Starting from sequence 1, we go upward in the reconstructed partition until we find the first internal node. We call this internal node *a.* All the sequences that descend from a, excluding the target sequence 1, are the genealogical nearest neighbors of sequence 1. The GNN proportion for the target sequence is the proportion of these neighbours that are case sequences within the clade below *a*. We repeat this calculation for all the target sequences and arrange the proportions in a vector indexed by target sequence. These proportions for the genomic region labeled as *A* comprise a vector, *G_A_*. For example, in partition *A,* the GNN proportion for the four target sequences are: 1/1 = 1 for sequence 1, 1/1 = 1 for sequence 2, 2/2 = 1 for sequence 3, and 2/3 = 0.67 for sequence 4. Thus *G_A_* = [1, 1, 1, 0.67]. Similarly in partition B we obtain *G_B_* = (1, 1, 0.67, 1).

Once we compute *G_A_* and *G_B_*, these vectors are weighted by the proportion of genomic region spanned by their respective partitions. Since partition A spans 60% of the total region of 10 kbp, the corresponding weight, *W_A_* = 0.6. Similarly *W_B_* = 0.4.

We are now ready to compute the average GNN proportion by taking the weighted average of all these proportions in both partitions. By taking the weighted average, we assign more weight to the partitions corresponding to long physical lengths of sequence than partitions corresponding to short physical lengths of sequence. This weighted average summarizes the proportion of nearest neighbours to the target sequence that are case sequences. In our example, the average GNN proportion can be computed as:

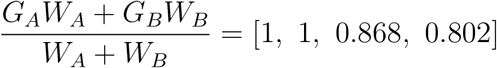

**Figure A.3:**
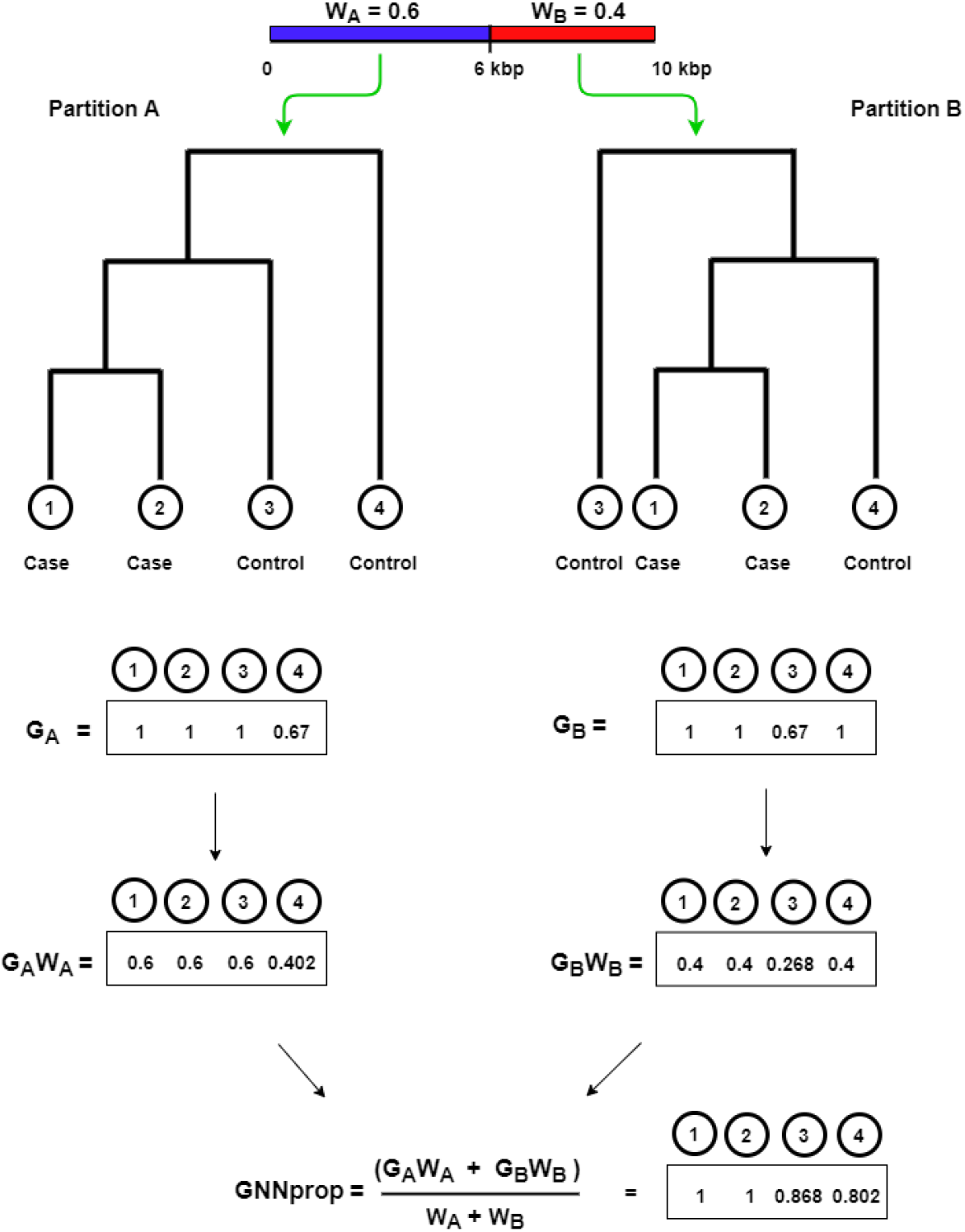
A worked example to illustrate the calculation of average GNN proportion. Four sequences are considered, labeled with circles 1 to 4, over a 10 kbp region.

### A.5 The estimated type-I error rate

**Table A.1:** The estimated type-I error rate or proportion of 500 null datasets that incorrectly reject the null hypothesis 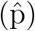 and associated approximate 95% confidence interval. Four methods are compared: 1) Fisher’s exact test (FET), 2) SKAT-O, 3) distance correlation (dCor), and 4) Mantel.

| Method | $\hat{p}$ | Approximate 95% CI | |
| --- | --- | --- | --- |
|  |  | Lower bound | Upper bound |
| FET | 0.040 | 0.022 | 0.057 |
| SKAT-O | 0.042 | 0.024 | 0.060 |
| dCor | 0.038 | 0.021 | 0.055 |
| Mantel | 0.048 | 0.029 | 0.067 |

